# Real-time Imaging of Decompression Gas Bubble Growth in the Spinal Cord of Live Rats

**DOI:** 10.1101/2024.02.09.578916

**Authors:** Roman Alvarado, Ulrich M. Scheven, J.-C. Meiners

## Abstract

1)

**Purpose:** To observe the growth and resolution of decompression gas bubbles in the spinal cord of live rats in real time using magnetic resonance imaging (MRI).

**Methods:** We constructed an MRI-compatible pressure chamber system to visualize gas bubble dynamics in deep tissues in real time. The system pressurizes and depressurizes rodents inside an MRI scanner and monitors their respiratory rate, heart rate, and body temperature while providing gaseous anesthesia under pressure during the experiments.

**Results:** We observed the formation of decompression gas bubbles in the spinal cord of rats after compression to 7.1 bar absolute and rapid decompression inside the MRI scanner while maintaining continuous gaseous anesthesia and vital monitoring.

**Conclusion:** We have shown the direct observation of decompression gas bubble formation in real time by MRI in live, anesthetized rats.

## 2) Introduction

Decompression illness is broad spectrum of clinical symptoms that arise from a rapid drop in ambient pressure, most notably in sports and commercial diving,^1^ caisson construction, and high-altitude aerospace operations.^2^ Among the various forms of decompression illness, spinal cord decompression sickness (SC-DCS) is particularly severe, resulting in paresthesia, sensory and motor deficits, and in some cases, death.^3^

Decompression sickness is generally thought to be caused by the formation of inert gas bubbles in the body that arise from a supersaturation of the tissue with gas upon a rapid reduction in the ambient pressure.^3^ Aural Doppler ultrasound and two-dimensional echocardiography have long been used to monitor the formation of circulating decompression bubbles in large veins or the heart.^4^ However, the correlation between the observation of these large vascular bubbles and clinical symptoms and markers of DCS, such as complement activation, is generally poor,^5^ as most damage arises from bubbles in deeper tissues rather than those circulating in the larger vasculature. Gas bubbles in the spinal cord are, up to now, only examined postmortem using pathology, which is rather limiting as it reveals no dynamic information, and it is fraught with artifacts from fixation and slicing that are often hard to distinguish from gas bubbles.^6^

We demonstrate that it is possible to observe the formation of gas bubbles in deeper tissues directly *in-vivo* using MRI. However, there are considerable technical challenges in combining the hyperbaric environment with MRI and maintaining anesthesia under pressure. In this proof of concept note, we present a hyperbaric chamber for small animals that is used inside an MRI scanner (N=2). It can maintain gaseous anesthesia in the animal throughout the pressurization and decompression experiment, and it allows monitoring of breathing, heart rate, and temperature. Additionally, it has a heat exchanger to warm the breathing gas mixture just before entering the chamber to maintain the animal’s body temperature. We present MRI data taken with this setup that show the formation of gas bubbles in the spinal cord of one of two rats in real-time upon decompression from 7.1 bar absolute to normobaric conditions.

## 3) Methods

### Magnetic Resonance Imaging and Pressure Protocol

Images were acquired in a 7T pre-clinical scanner (Varian, now Agilent), using a 60mm saddle coil manufactured by Morris Instruments Inc. Four-to six-week-old female Sprague Dawley rats were imaged inside a custom-built MRI-compatible pressure chamber. A standard gradient echo multi-slice (GEMS) sequence was employed (TR/TE = 500ms/2.5ms, FA=30, voxel size 125x125x1000 μm3). For this exploratory work we chose a gradient echo sequence because the images are sensitive to both spin density – the absence of spins inside a newly formed bubble – and the field distortions caused by the susceptibility contrast between the empty bubble and the tissue surrounding it.

With the rat inside the pressure chamber, and the chamber placed in the scanner, the chamber was pressurized to 7.1 bar absolute in 45 s. The pressure was maintained for 20 min, and then decompressed to 1 bar absolute within 30 s. This protocol is known to preferentially induce neurologic DCS in rats.^7^ We show several MRI scans taken from 8 minutes into the compression phase, to 10 minutes after decompression.

### Image Processing

The MRI images were processed using OpenCV algorithms programmed in Python. To enhance image quality, each image underwent a two-fold, cubic interpolation, upsampling in both dimensions. A multi-nonlocal means denoising algorithm was then applied using the two temporally closest scans. This entails using a small patch of pixels in the image of interest and comparing the same patch of pixels in the time forward image and time backward image. The weighted average of these pixels replaces the pixel in the image of interest and thus, reduces noise. This process is repeated for every pixel in each image, for all images. Further noise reduction and feature edge enhancement were achieved through the implementation of a five-pixel diameter bilateral filter. Lastly, image contrast was improved using a contrast-limited adaptive histogram equalization algorithm on each image where the histogram intensities of small patches of pixels were equalized or uniformly distributed if they were above a specified pixel intensity threshold.

### Chamber and pressurization system

The cylindrical pressure chamber is machined from solid, transparent polycarbonate. The tube has a total length of 330 mm, an outer diameter of 5.74 mm, and a wall thickness of 3.3 mm. An internal polycarbonate tray supports the rat laid in a supine position. The pressure chamber tube has threads and O-ring seals on both ends to attach an end cap with electric and gas feedthroughs on one end, and a heat exchanger for the inflowing gas on the other end. The electrical feedthroughs connect to the ECG and temperature sensors. A 5/32” OD push-to-connect fitting connects to a pressure sensor (Dwyer Instruments 626-04-GH-P8-E2-S5) to instantaneously monitor the chamber pressure. A second fitting serves as the exit port for the anesthetic gas flow. (Figure 1).

**Figure 1:**
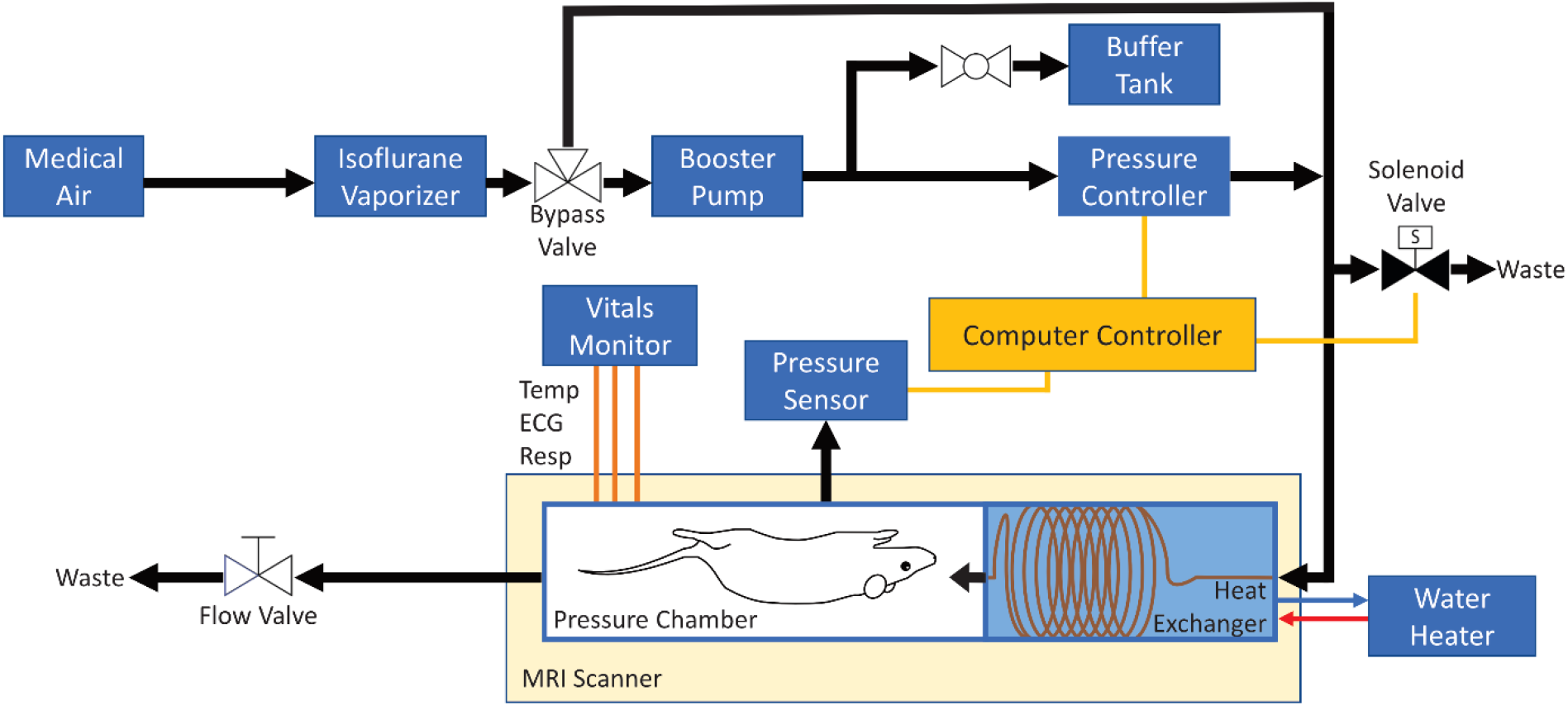
Schematic of the chamber and pressurization system. The air pathway is indicated by black arrows, electronic connections are indicated by yellow lines, vital monitoring wiring is indicated by orange lines, and hot water is indicated by a red arrow, while cold water is indicated by a blue arrow.

Pressure inside the pressure chamber is controlled with an electronic pressure control system. Medical-grade air mixed with isoflurane, is compressed by a booster pump (Haskel AGD-4) to pressurize medical grade air mixed with isoflurane to 9.3 bar absolute and stored at that pressure in a small buffer tank (Figure 1). An electronic pressure-control valve (ProportionAir QPV1MANEEZP100PSGBXN) is used in conjunction with the chamber pressure sensor and custom LABVIEW software to implement the desired pressure profiles in the chamber. An additional solenoid valve (Atlantic Valves BZW-20) is used to quickly depressurize the system, as depressurization through the control valve itself is too slow for our protocol.

#### Anesthesia and vital monitoring

Since the anesthetic effect of isoflurane depends primarily on the partial pressure rather than its absolute concentration,^8^ we administered an isoflurane concentration of 0.25% (v/v) to achieve stable anesthesia at 7.1 bar absolute. To simplify anesthetic maintenance at atmospheric pressure before compression and after decompression, we used a bypass valve to circumvent the booster pump and pressure control equipment during these phases of the experiment. This allowed us to directly feed the output of the evaporator into the pressure chamber at 1% (v/v) isoflurane concentration.

The pressure cell was designed to accommodate commercial MRI-compatible vital monitoring hardware from Small Animal Instruments, Inc. This monitoring suite includes the Model 1030 Monitoring and Gating Systems, 3-lead surface ECG electrodes, and a rectal thermistor to monitor the animal’s temperature. Respiration is monitored using a technique termed E-RESP by the SA-Instruments: the ECG wires are placed to make a loop around the animal’s chest (Figure 2). As the loop size changes with respiratory motion, a small emf voltage is induced in the ECG wires, due to the changing magnetic flux through the loop. We employed this E-RESP method of respiration monitoring because conventional pneumatic pillows are unsuitable for operation at elevated pressures. Vitals were measured and recorded using the software provided by Small Animal Instruments.

**Figure 2:**
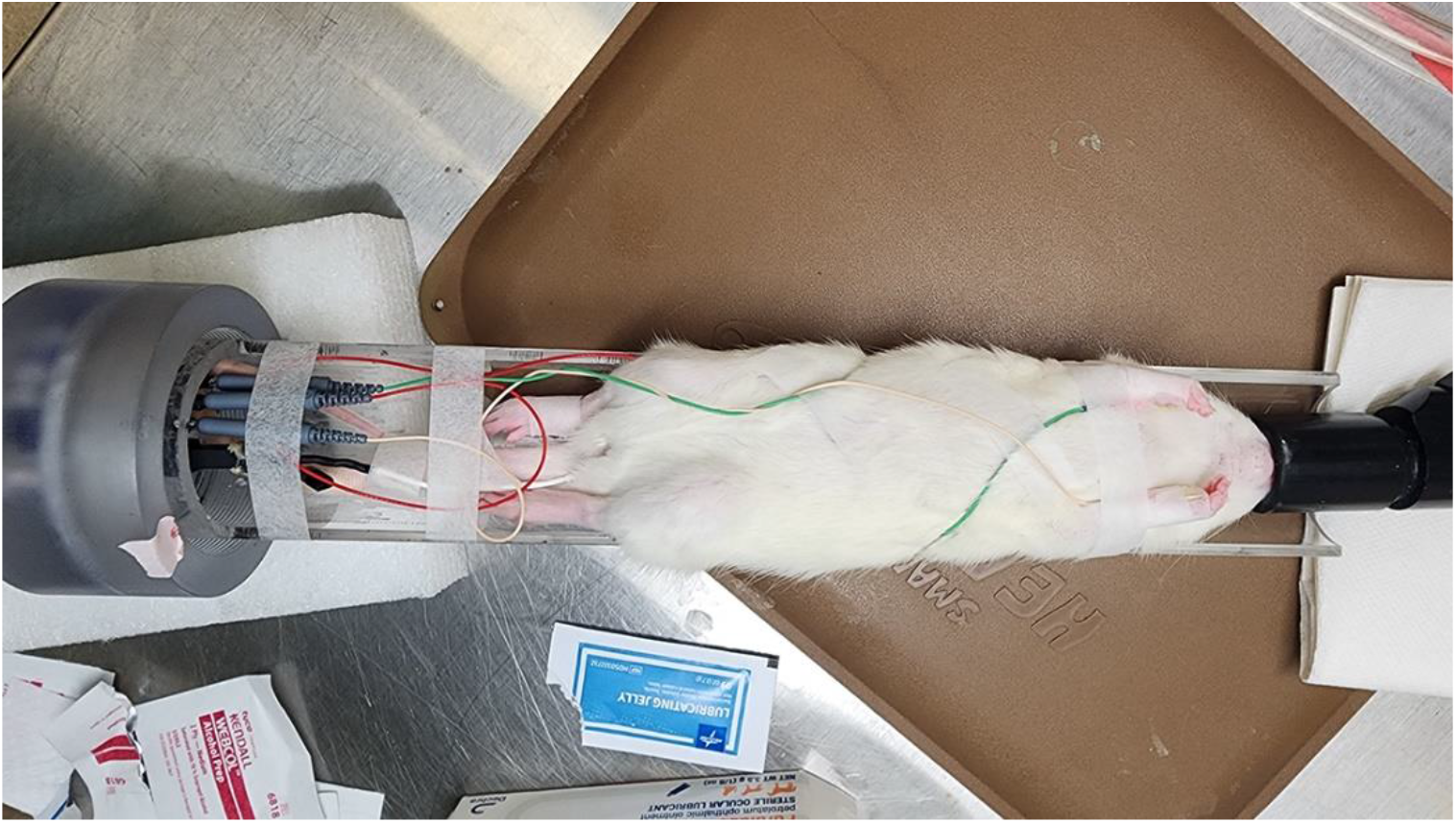
A rat is secured on the pressure chamber tray with ECG wires configured for E-RESP and an inserted rectal temperature sensor.

To compensate for heat loss in the animal during imaging, the breathing air can be warmed as it enters the chamber. To achieve this, we built an MRI-compatible heat exchanger that is mounted at the gas inlet end of the pressure chamber. The heat exchanger consists of a small compartment through which warm water is circulated. The breathing gas flows through a five-meter copper tubing coil that is in direct contact with the warm water. The temperature of the warm water can be manually adjusted to maintain the appropriate body temperature of the animal.

## 4) Results

We successfully compressed live rats inside the MRI scanner to 7.1 bar absolute for 25 minutes before decompressing them and maintaining them anaesthetized at atmospheric pressure until they succumbed to severe DCS. Real-time MRI scans during this process showed the formation of decompression bubbles in the spinal cord of the rat. Figure 3A -left, shows an axial image slice of the thoracic region in the compressed rat, before bubble formation. Figure 3A - right, taken approximately 10 minutes after decompression and around the time of the rat’s expiration, displays a large gas bubble in the posterior median spinal vein. A smaller gas bubble in the nervous tissue of the spinal cord is also visible. We further note distortions in the anterior region of the spinal cord but cannot conclusively associate them with anatomical features such as posterior spinal veins or the meninges.

**Figure 3:**
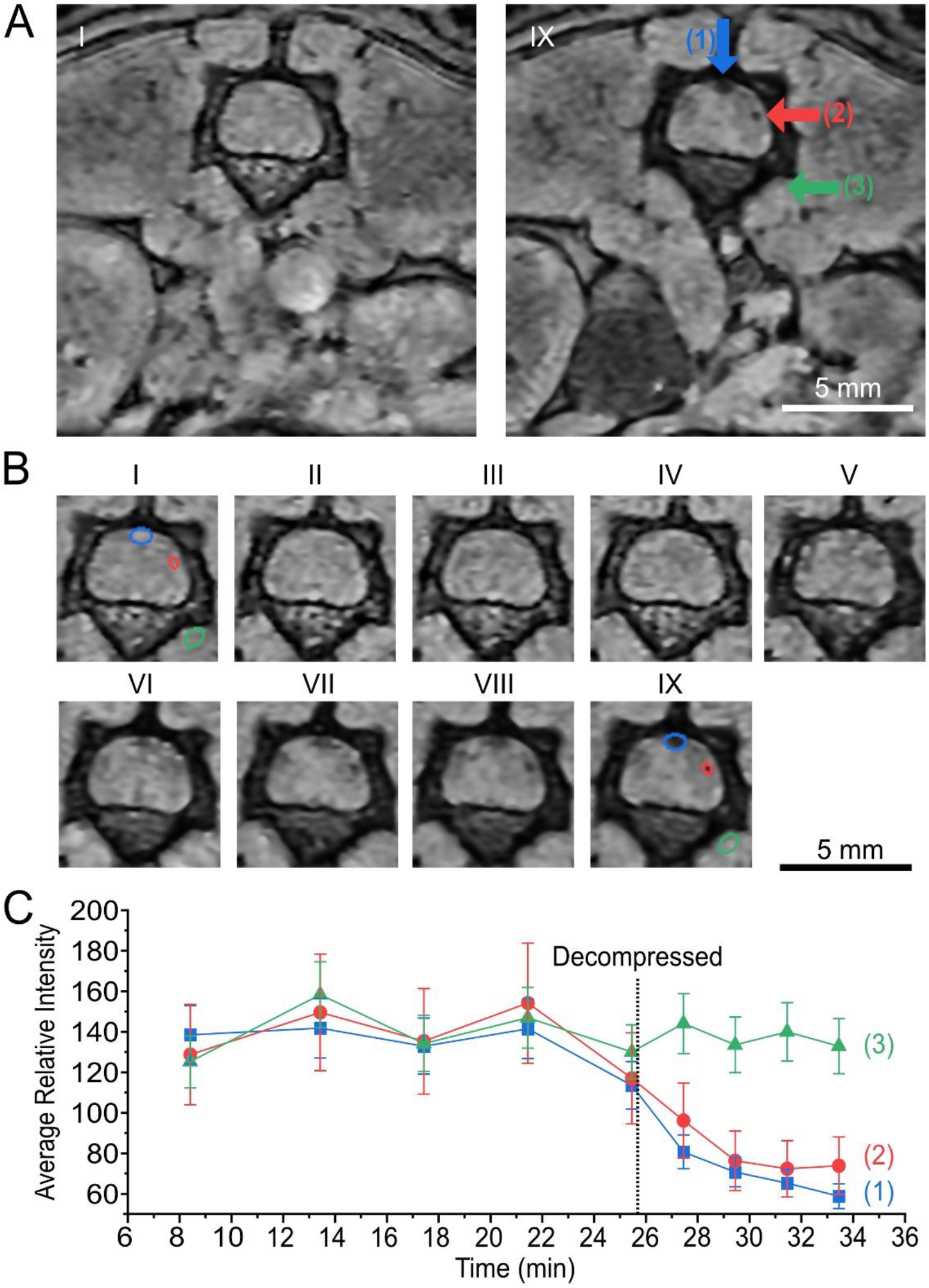
MR images of the spinal region of a rat (A) (left) after pressurization to 7.1 bar absolute and (right) 10 minutes after decompression to 1 bar absolute. The second image shows a gas bubble in the posterior median spinal vein (vertical blue arrow), one other suspected gas bubble (horizontal red arrow), and a non-bubble control region (horizontal green arrow). The right third of the spinal cord is darker in (right), possibly indicating hemorrhage here. (B) A time series of MRI scans of the spinal cord taken during the experiment are shown. An ellipse region of interest, one for each bubble and the control region, were used to determine the (C) average intensity over the series of images to determine the appearance of bubble formation. The intensity timepoints indicate the start of the MRI scan. The bubble in the posterior median spinal vein is shown in blue (1), the other suspected bubble is shown in red (2), a non-bubble control region is shown in green (3).

The evolution of bubble formation is shown in Figure 3B. The images while the chamber was compressed (I-IV) showed no significant change while images post decompression (VI-IX), revealed bubbles. Images VI-VIII showed potential lateral vein fracturing occurring which may be responsible for the appearance of hemorrhaging near the lateral most bubble. An ellipse region of interest is shown in Figure 3B-1 and 3B-IX to determine the change in average intensity. As indicated in Figure 3C, the average intensity decreases with the formation of a bubble in the posterior median spinal vein and in the lateral most bubble, but not in the non-bubble control region.

Adequate anesthetic depth was successfully maintained throughout the experiment. Continuous monitoring of respiration rate, heart rate, and temperature was conducted during pressure changes, as depicted in Figure 4. The heart rate stayed around 300 beats per minute and the respiratory rate was approximately 30-35 breaths per minute until the rat expired.

**Figure 4:**
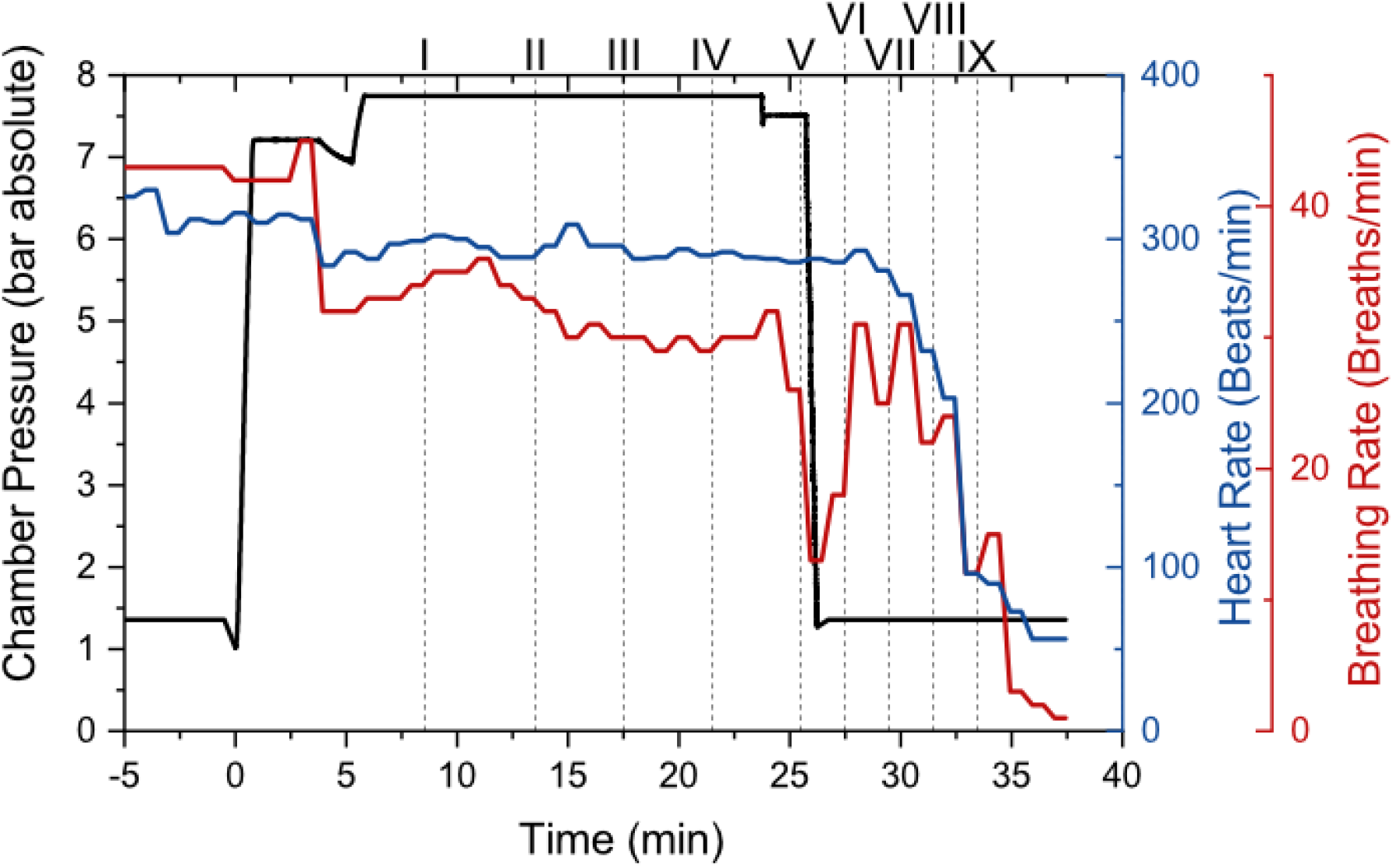
Heart rate and respiratory rate of the rat during compression and decompression. The vertical dashed lines indicate the times at which MRI scans were taken and correspond to those shown in Figure 3.

## 5) Discussion and Conclusions

Decompression sickness symptoms are closely related to the formation of inert gas bubbles in the body.^3^ The exact pathophysiology of how gas bubbles in deeper tissues form, and subsequently dissolve upon treatment through recompression remains largely mysterious, as gas bubbles in the spinal cord and similar tissues could only be observed through histopathology until now. Our novel combined pressurization and imaging method allows the observation of decompression gas bubble formation in a live animal under anesthesia in a pressure chamber inside the MRI scanner in real time. We demonstrate a 125 μm voxel resolution for a stack of 30 3.2 cm x 1.6 cm images. This resolution is achieved with four averages for a four-minute scan pre-decompression and two averages for a two-minute scan post-decompression.

We observed the formation of a large bubble with a cross section of approximately 0.4 mm^2^ in the largest vein of the spine, the posterior median spinal vein, and a smaller bubble in another area of the spinal cord. The physical deformations and distortions of the tissue surrounding the bubbles in response to their rapid growth may have led to direct mechanical damage of the tissue and local hemorrhage, eventually leading to the demise of the animal.

In conclusion, we have shown that it is possible to observe the growth of decompression gas bubbles in real time in live rats using MRI in conjunction with a purpose-built pressure cell and life support and monitoring equipment. Using an aggressive compression and decompression protocol that is known to preferentially induce SC-DCS, we were able to observe the formation of gas bubbles in the spinal cord until the animal expired from the damage. Maintenance and monitoring of anesthesia were possible at isoflurane concentrations that are largely determined by the partial pressure of the anesthetic during the various stages of the experiment. This new capability will enable rigorous testing of many of the assumptions that are made in understanding the pathophysiology of deep-tissue DCS and its treatment though recompression without having to rely solely on histopathology.

## 6) Acknowledgements

This study was funded by the Divers Alert Network. We thank Angela Yee for her assistance in handling animals during the experiments. This work was carried out with the approval of the Institutional Animal Care and Use Committee (IACUC). All data and imaging processing scripts are available upon request.

